# Dual Mechanism of Action: Exosomes from Human iPSC-Cardiomyocytes and Mesenchymal Stem Cells Restore Injured Myocardium

**DOI:** 10.1101/2024.05.29.596548

**Authors:** Eileen Tzng, Nathan Bayardo, Gentaro Ikeda, Hiroyuki Takashima, Jennifer Lyons, Mihoko Bennett, Connor G O’Brien, Phillip C. Yang

## Abstract

**Background:** Transplantation of mesenchymal stem cells or induced pluripotent stem cell derived cardiomyocytes improve heart function after myocardial infarction in pre-clinical models. Exosomes are extracellular vesicles, 30-150nm in size, which regulate the paracrine signal of the stem cells. We investigated the functional outcomes and biological effects of exosomes from pure populations of human bone marrow derived mesenchymal stem cells (MSCs) and induced pluripotent stem cell derived cardiomyocytes (iCMs) in a porcine acute myocardial infarction model.

**Methods:** Yorkshire swine were subject to proximal left anterior descending artery occlusion with a catheter balloon for 1 hour for ischemia-reperfusion injury. Ten 500ul injections containing 5 x 10^11^ exosomes isolated from the tissue culture media of iCMs or MSCs were delivered transendocardially into the peri-infarct region. Cardiac function was assessed by magnetic resonance imaging (MRI). Multi-omic analyses were performed in the *ex vivo* swine peri-infarct specimen to delineate the mechanism of action.

**Results:** Cardiac MRI at weeks 2 and 4 showed significant improvement in heart function in iCM-derived exosomes while MSC-derived exosomes showed a trend towards improvement. A comparative analysis of transcriptomic sequencing of the porcine peri-infarct tissue and Next Generation Sequencing of the exosome cargo confirmed the dual mechanism of action. The marked improvements seen in cardiac function are conferred by miRNA carried by the exosomes, particularly by cardioprotective reduction in metabolism during acute myocardial injury while promoting concurrent cardiomyocyte cell cycle re-entry and proliferation.

**Conclusions:** Significant reduction in myocardial metabolism and increase in proliferation signal pathways were found in both exosome treatment groups; however, distinct sets of microRNAs were found to underlie the mechanism of action in each population of exosomes.

## Introduction

Transplantation of mesenchymal stem cells (MSCs) or induced pluripotent stem cell derived cardiomyocytes (iCMs) improve heart function after myocardial infarction^1–3^. Acute myocardial infarction restricts the perfusion to the myocardium, producing irreparable damage to the left ventricle and contributing significantly to the prevalence of heart failure. Today, heart failure is the leading cause of hospital admission in the United States with continued increase in patient population due to aging and chronic diseases such as diabetes and obesity^4^. The inability of the adult heart to regenerate the damaged myocardium contributes to scarring, significantly attenuating cardiac contractility and, subsequently, left ventricular ejection fraction (LVEF). The resultant scar tissue contributes to the downward spiral of heart failure as the collagen deposits do not contribute to cellular bioenergetics or tissue homeostasis, introducing metabolic maladaptation to compensate for increased bioenergetic needs.

The application of MSCs, induced pluripotent stem cells, and cardiosphere derived cells into the injured heart has improved cardiac function in multiple pre-clinical models^1, 2, 5–6^. Yet, cell engraftment in the myocardium presents a multitude of biological challenges such as immune rejection, survival, and ventricular arrhythmia. The therapeutic effects of cardiospheroid cells vs. small extracellular vesicles derived from cardiospheroid cells in ischemic swine hearts demonstrated superior efficacy of the cardiospheroid cell-derived extracellular vesicles^7^. The extracellular vesicles obviated the need for cell engraftment and eliminated arrhythmia for their therapeutic efficacy. Identification of specific effectors and mechanisms responsible for the therapeutic effects is complex due to the heterogeneous population of extracellular secretomes. Exosomes are a subclass of extracellular vesicles, 30-150nm in size, characterized by the integral membrane proteins CD81, CD63 and CD9. These small vesicles are secreted by most cell types and deliver protein, miRNA, mRNA, and other biologically active molecules to neighboring or recipient cells. Exosome cargo is determined by the transcriptional state or identity of the donor cell with different cell types eliciting different paracrine effectors.

Though effects of exosome administration on heart function following myocardial infarction have been demonstrated in countless models, the specific mechanisms of the biologically active cargo remain poorly characterized^8–11^. Exosomes contribute to a wide spectrum of signaling processes both pathologic and physiologic. Exosomes may play a role in cancer metastasis through activation of oncogenic pathways while also maintaining cardiac tissue homeostasis. Defining the mechanisms of exosome signal remains elusive given that cargo loading is contingent on the transcriptional state of the donor cells, different cell types, and physiological stimuli, yielding exosomes with unique cargos.

Brief periods of ischemia augment exosome cargo loading and release. Ischemic conditions *in vitro* modulate exosomal cargo in iCMs through upregulation of miR Cluster 106a-363^11^. Ischemic preconditioning results in reduction of opening of mitochondrial permeability transition pore and reactive oxygen species production in the injured myocardium^12^. Subsequently, plasma exosome concentrations increase, conferring cardio protection through reducing cell death and/or mammalian target of rapamycin (MTOR)^13–15^. Our group has previously shown that exosomes from iCMs are cardioprotective in a mouse left anterior descending artery ligation model through induction of mitophagy by inhibition of the MTOR pathway, which is argued to be a necessary event during ischemic preconditioning^16^. Furthermore, cardiac metabolism has been modified to improve outcomes in the failing heart, specifically using SGLT2 inhibitors and other antidiabetic drugs^17^. Therefore, there is an intrinsic link between ischemic preconditioning, metabolic augmentation, and exosome signaling.

To investigate this link and elucidate the mechanism of action, we compared the effects of the exosomes derived from MSCs (MSC-Exo) or iCMs (iCM-Exo) in this study. The exosomes were delivered transendocardially in a porcine ischemia reperfusion injury model. Transcriptomic analysis of the porcine peri-infarct myocardium, which is most vulnerable to remodeling and/or electrical abnormality, treated with either iCM-Exo, MSC-Exo, or PBS revealed a concerted reduction of myocardial metabolism in both treatment groups. Correlation between tissue transcriptome and Next Generation Sequencing of the exosomes was then used to study the biological effectors within the MSC-Exo vs. iCM-Exo.

## Methods

Animal care and interventions were done in accordance with the Laboratory Animal Welfare Act and Stanford University Administrative Panel on Laboratory Animal Care. All animals received humane care and treatment in accordance with the “Guide for the Care and Use of Laboratory Animals” (www.nap.edu/catalog/5140.html). Studies were performed under ARRIVE guidelines as endorsed by Stanford University. Studies were performed under APLAC protocol #16066. All research was performed at Stanford University in Stanford, CA. The data that support the findings of this study are available from the corresponding author upon reasonable request.

### Bone Marrow Mesenchymal Stem Cell

StemPro™ BM Mesenchymal Stem Cells (ThermoFisher Scientific) were cultured in MesenPro Media (12746012, ThermoFisher Scientific) in flat surfaced 75cm^2^ Tissue Culture Flasks. For media collection, MSCs at 60% confluence were washed twice with PBS to remove MesenPro serum and incubated in high glucose DMEM with 10% exosome free FBS for 48 hours.

### iPSC-Derived Cardiomyocytes

Induced pluripotent stem cells were cultured with Essential 8 pluripotent stem cell medium (E-8 media, Life Technologies, CA) in Matrigel (Corning, NY) coated 6 well plates. Differentiation was initiated at 80% confluence using GSK inhibitor (6uM CHIR) in basal iCM differentiation medium (Gibco® RPMI 1640 medium, GlutaMAX™ Supplement, ThermoFisher Scientific) with B-27 Supplement Minus Insulin (A1895601, ThermoFisher Scientific) for day 0-2. Media was replaced with Gibco® RPMI 1640 medium with B-27 Supplement Minus Insulin on day 3. Reprogrammed iPSCs were then incubated in 2uM C59 (WNT inhibitor) in basal iCM differentiation medium with B-27 Supplement Minus Insulin for days 4-6. Cells were changed into basal iCM differentiation medium with B-27 Supplement Minus Insulin until contractility was observed around day 10-14. Cells were then incubated in iCM maintenance media (Gibco® RPMI 1640 medium, GlutaMAX™ Supplement, ThermoFisher Scientific) with B-27 Supplement for (17504044, ThermoFisher Scientific) 48 hours for purification and replated onto Matrigel coated plates. iCMs were allowed to mature for 7 days before collecting supernatants. The purity of iCMs was assessed with flow cytometry. iCMs were cultured in iCM maintenance media (Gibco® RPMI 1640 medium, GlutaMAX™ Supplement, ThermoFisher Scientific) with B-27 Supplement for 2 days before harvesting media for exosome isolation.

### Exosome Isolation and Characterization

Exosomes were isolated according to previous validated protocols^9–11^. iCMs and MSCs were washed with PBS and incubated in exosome free media for two days. Supernatant was collected and centrifuged at >1000g for 10 minutes, passed through a 0.22-μm syringe filter, and incubated with 25% polyethylene glycol 8000 (PEG-8000, 25322-68-3, Sigma Aldrich, MO) at 4°C overnight. The resulting mixture was then centrifuged at 1500g for 30 minutes and aspirated to leave 5ml over the pellet. This was centrifuged again at 300g for 5 minutes, the remaining supernatant was removed, and the pellet was resuspended in filtered PBS and stored in -80°C.

### Nanoparticle Tracking Analysis (NTA)

Size and concentration of exosomes were evaluated by NTA with a NanoSight LM20 (NanoSight, UK). Samples were loaded into the sample chamber with sterile syringes and imaged using a 640 nm laser. Three measurements of the same sample were performed. Particle size and velocity in the fixed chamber was used in proprietary software (NTA 3.1 Build 3.4.4, NanoSight Ltd) to determine the mean size and +SD values of the particles. Mean and standard deviation were calculated as per software recommended protocol. NTA was used to reveal the concentration of particles in exosome pellets to aliquot the desired injection concentration.

### Porcine Ischemia Reperfusion Model

Animal care and interventions were done in accordance with the Laboratory Animal Welfare Act and Stanford University Administrative Panel on Laboratory Animal Care. All animals received humane care and treatment in accordance with the “Guide for the Care and Use of Laboratory Animals” (www.nap.edu/catalog/5140.html). Studies were performed under ARRIVE guidelines as endorsed by Stanford University. Studies were performed under APLAC protocol #16066. With approval and under the guidelines of Stanford Animal Care and with Stanford Veterinary Service Center (VCS), 14 female Yorkshire swine (age 3-5 months for duration of model) were anesthetized with 1-2% isoflurane and subject to myocardial infarction using a 10-mm over-the-wire angioplasty balloon inserted through the femoral artery and placed in the proximal left anterior descending coronary artery (LAD) at the first diagonal branch and inflated for 60 min and, subsequently reperfused. Occlusion of the LAD was confirmed by contrast agent using the Allura Bi-plane Fluoroscopy unit (Philips, Netherlands). Immediately following deflation, a Biocardia Helix-Morph catheter was used for transendocardial delivery of ten 500 µl injections of resuspended exosomes, administering a total of 5x10^11^ particles to the peri-infarct region. Interventionalist and MRI tech were blinded to the treatment group.

### Intramyocardial Injection

Swine were administered daily cyclosporine A (10 mg/kg oral administration, Sandimmune formulation, Novartis Pharma AG) for three days before and two days after the point of delivery. At cell delivery, a 1-time dose of 15 mg/kg methylprednisolone sodium succinate was administered intravenously. Ten 0.5 mL injections containing PBS (n=4) or 5 x 10^11^ exosomes isolated from either iCMs (n=5) or MSCs (n=5) were delivered transendocardially to the peri-infarct region using Helix Helical Infusion System, or a Morph Universal Deflectable Guide Catheter (BioCardia Inc, CA) as previously described^5^. The Allura Bi-plane Fluoroscopy unit (Philips, Netherlands) was used to visually administer injections and map injection sites around the infarction site.

### Magnetic Resonance Imaging

MRI was acquired at week 2 and week 4 after treatment to evaluate heart function and viability of the peri-infarct region as described previously^18^. Cardiac MRI was performed using Signa 3T EXCITE scanner (GE Healthcare) and a phased array 2 channel surface coil (Rapid MR international). EVP 1001-1 (Eagle Vision Pharmaceutical Corp) was administered intravenously at a dose of 0.7 mL/kg for in vivo manganese enhanced MRI (MEMRI). Right and left anterior-oblique ventriculograms were generated to identify the peri infarct region with MEMRI and delayed gadolinium enhancement magnetic resonance imaging (DEMRI). Briefly, cine images were acquired using steady state free procession (TR 3.4 ms; TE min-full; flip angle 45°; thickness 8 mm; matrix 224 × 224; and field of view [FOV] 28 cm). MEMRI was obtained using fast gradient echo-inversion recovery (FGRE-IR) sequence (TR 6.2 ms; TE 2.9 ms; flip angle 15°; thickness 8 mm; matrix 224 × 192; FOV 28 cm; TI 300–600 ms) 25–40 min after IV infusion of EVP1001-1 at 20.3 μmol/kg (2.0 μmol/kg/min) (Eagle Vision Pharmaceutical Corp; Downington, PA). After a 30-minute washout period, 0.2 mmol/kg of Gadolinium-DTPA (Magnevist, Bayer Health Care Pharma AG, Berlin, Germany) was administered. DEMRI was obtained using FGRE-IR sequence (TI 250–300 ms) 10–25 min later.

### Post-Mortem Tissue Staining and Harvest of Peri-Infarct Tissue

At week 4 following myocardial injury, the hearts were perfused with Evans Blue, ex-planted, and stained with tetrazolim chloride (TTC) to delineate the peri-infarct region^19^. 20 to 30 ml of 1% TTC solution was injected through the guidewire lumen of the over-the-wire balloon catheter, which stains viable myocardium containing mitochondrial dehydrogenase enzymes red and leaves non-viable myocardium white (infarct core). 1% Evans blue dye was injected into the left (60 ml) and right (30 ml) coronary arteries to delineate Evans blue negative area at risk and Evans blue positive perfused (normal) myocardium. Finally, the pig was euthanized with an intravenous potassium chloride injection. The excised heart was cut into 1 cm-thick cross-sectional slices and fixed with 10% formalin. The slices were cut into pieces containing peri-infarct and remote zones and embedded in paraffin. The sliced sections were stained with hematoxylin eosin (H&E) for morphological observation and picrosirius Red to visualize myocardial scarring. Tissue was harvested from viable myocardium (TTC positive) with no perfusion (Evans-blue negative) and RNA was isolated and subsequently sequenced with Illumina Novaseq 6000. Reads were aligned to the pig genome and Deseq2 was used to perform differential expression analysis in the peri-infarct tissue treated with either exosomes or PBS.

### RNA Isolation

Total RNA from harvested pig tissue was isolated using the Qiagen Rneasy Plus Minikit (74134, Qiagen, Germany), following the supplier recommended protocol.

### Tissue RNA Sequencing

RNA samples were submitted to Novogene (CA) for RNA-sequencing on Illumina Novaseq 6000, S4 flowcell. In brief, a total amount of 1 µg RNA per sample was used as input material for the RNA sample preparations and sequencing libraries were generated using NEBNext® Ultra TM RNA Library Prep Kit for Illumina® (NEB, USA).

### Exosomal RNA Sequencing

Conditioned media (n=2 each group) were sent for Exo-NGS™ services (System Biosciences, CA, USA). In brief, extracellular vesicles were isolated using SBI’s ExoQuick® Exosome Isolation kit protocol and SBI’s spin column-based RNA Purification Kit. Next Generation Sequencing was run on Illumina Nextseq at an approximate depth of 10-15 million reads per sample.

### Bioinformatic Analysis

Sequencing analysis of exosomal RNA-sequencing was performed by Banana Slug Genomics Center at University of California Santa Cruz using Exosome Small RNA-seq Analysis 1.10.

Images and graphs of analysis of tissue RNA-seq were provided with analysis. Briefly, Illumina sequencing was used and trimmed reads were then mapped to the human genome. Raw read counts were calculated for known gene categories including ncRNAs, antisense transcripts, coding and intronic regions of mRNAs, and repeats. Raw read counts were normalized across all samples and differential expression analysis was performed using DEseq. Significant differentially expressed genes were determined by adjusted P-value with a threshold of 0.05. Refer to documentation provided by Novogene for additional information.

Significant DEGs were subsequently analyzed by Enrichr^20–22^ for GO and KEGG analysis using the top 100 most significant genes for pig tissue RNA-seq analysis and using all significant genes for mRNA RNA-seq analysis of exosome cargo. Graphs of DEGs were created in and obtained from Appyter^23^. Heatmap of significant differentially expressed miRNAs was created in R Statistical Software (v 4.3.1; R Core Team 2023) using DEG lists generated by Banana Slug Genomics Center at University of California Santa Cruz.

### Statistical analysis

Data are expressed as median with interquartile range. Differences among 3 groups or more were assessed using Kruskal Wallis test followed by Dunn’s multiple comparisons to account for small sample size on GraphPad Prism version 10.1.2 for Windows, GraphPad Software, Boston, Massachusetts USA, www.graphpad.com. To determine whether Kruskal Wallis can be used, normality, homoscedasticity, and other assumptions were determined using residual plot, homoscedasticity plot, and QQ plot. p values <0.05 were considered statistically significant.

## Results

### Exosomes from iCMs and MSCs preserved cardiac function after ischemia reperfusion injury

Porcine hearts were treated with iCM-derived exosomes (iCM-Exo, n=5), MSC-derived exosomes (MSC-Exo, n=5), or PBS (n=4). Cardiac MRI at weeks 2 and 4 post-injury measured the following: left ventricular ejection fraction (LVEF), left ventricular end-systolic volume (LVESV), left ventricular end-diastolic volume (LVEDV), delayed enhanced gadolinium magnetic resonance imaging quantified the scar size, and manganese enhanced magnetic resonance imaging measured myocardial viability. LVEF increased significantly at week 2 by iCM-Exo vs. PBS (control) treatment groups (median (IQR); PBS 21.74 (18.67-23.82) vs iCM-Exo 31.80 (26.87-33.35)*, *p<0.05, Figure 1A). This significant improvement in LVEF at week 4 was sustained in the iCM-Exo treatment group (median (IQR); PBS 19.89 (14.19-21.91) vs iCM-Exo 33.05 (28.79-34.44)*, *p<0.05, Figure 1A). Furthermore, LVEF difference (LVEF delta) between weeks 2 and 4 were compared. LVEF delta showed significant improvement only by iCM-Exo vs. PBS (median (IQR); PBS -3.375 (-5.08-0.21) vs iCM-Exo 1.33 (0.85-2.125)*, *p<0.05, Figure 1A). Significant LVEDV (median (IQR); PBS 164.1 (141.3-206.9) vs iCM-Exo 109.8 (107.7-117.0)*, *p<0.05, Figure 1B) and LVESV (median (IQR); PBS 127.4 (109.8-167.0) vs iCM-Exo 74.92 (72.58-84.52)*, *p<0.05, Figure 1C) difference was only seen at week 2 in iCM-Exo vs. PBS. Finally, MSC-Exo vs. PBS treatment groups showed significantly decreased scar size only at week 4 (median (IQR); PBS 34.94 (33.83-37.38) vs MSC-Exo 31.39 (24.22-31.98)*, *p<0.05, Figure 1D). Cardiac MRI demonstrated that iCM-Exo treatment improved cardiac function.

**Figure 1.**
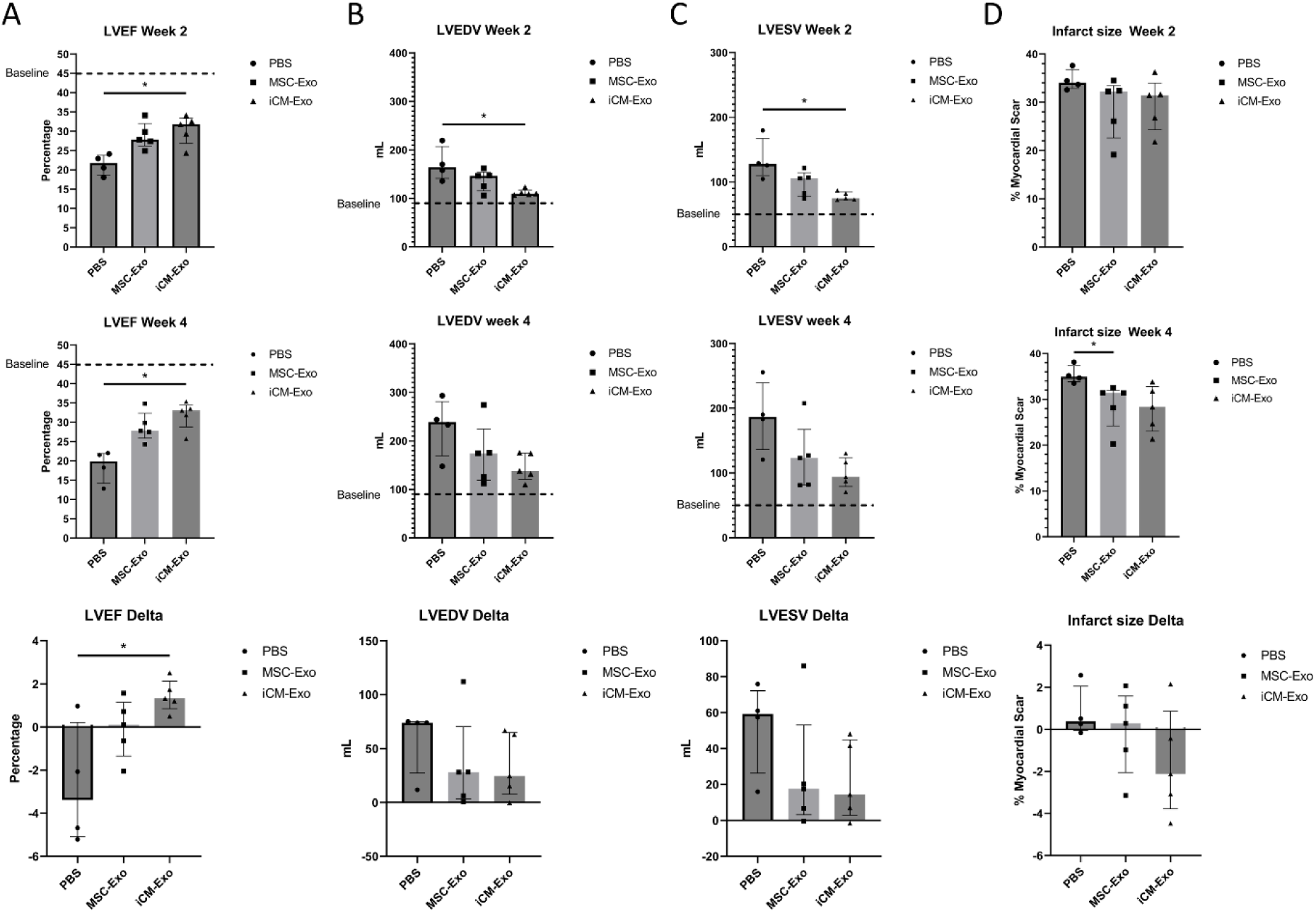
Cardiac MRI of porcine hearts treated with PBS, MSC-Exo or iCM-Exo. Quantification of (A) left ventricular ejection fraction, (B) left ventricular end diastolic volume, (C) left ventricular end systolic volume, and (D) infarct size as an area % of delayed gadolinium enhancement magnetic resonance imaging positive signal. Statistical significance was determined by Kruskal Wallis (iCM-Exo, n=5; MSC-Exo, n=5; PBS n=4) with multiple comparisons. Median with interquartile range is shown. Only significance between control and treatment groups within each week are shown (*p<0.05). iCM-Exo: exosomes derived from induced pluripotent stem cell derived cardiomyocytes; LVEDV: left ventricular end diastolic volume; LVEF: left ventricular ejection fraction; LVESV: left ventricular end systolic volume; MSC-Exo: exosomes derived from mesenchymal stem cells

### Exosomes from iCMs showed higher myocardial viability in the peri infarct region than MSC exosome treatment

At weeks 2 and 4, Yorkshire pigs were given gadolinium and manganese contrast agents to assess scar size and viable tissue, respectively. Manganese enhanced magnetic resonance imaging assessment showed that iCM-Exo treatment significantly increased myocardial viability at both week 2 (median (IQR); PBS 82.88 (77.72-83.73) vs iCM-Exo 89.52 (87.86-93.23)**, **p<0.01, Fig 2A) and week 4 (median (IQR); PBS 80.17 (78.17-83.23) vs iCM-Exo 92.48 (90.40-95.46)**, **p<0.01, Fig 2A). MSC-Exo treatment was not significant at either time points. Injections were performed into the peri-infarct region between scar tissue and healthy myocardium, suggesting that iCM-Exo provided significant rescue of the injured but viable cells in the peri-infarct region to improve myocardial function and morphology. While there was no significant improvement found, both treatment groups trended towards improvement in peri-infarct region viability compared to the control treatment (Fig 2B). Scar size measured by delayed gadolinium enhancement magnetic resonance imaging remained similar across all treatment groups due to the non-specific distribution of gadolinium. However, the viability specific manganese enhanced magnetic resonance imaging enabled significant detection of enhanced myocardial viability by the treatment group. iCM-Exo treatment restored the peri-infarct region and improved the cardiac function, volumetric remodeling and morphology in porcine chronic heart failure model at week 2 (Fig 1B, 1C).

**Figure 2.**
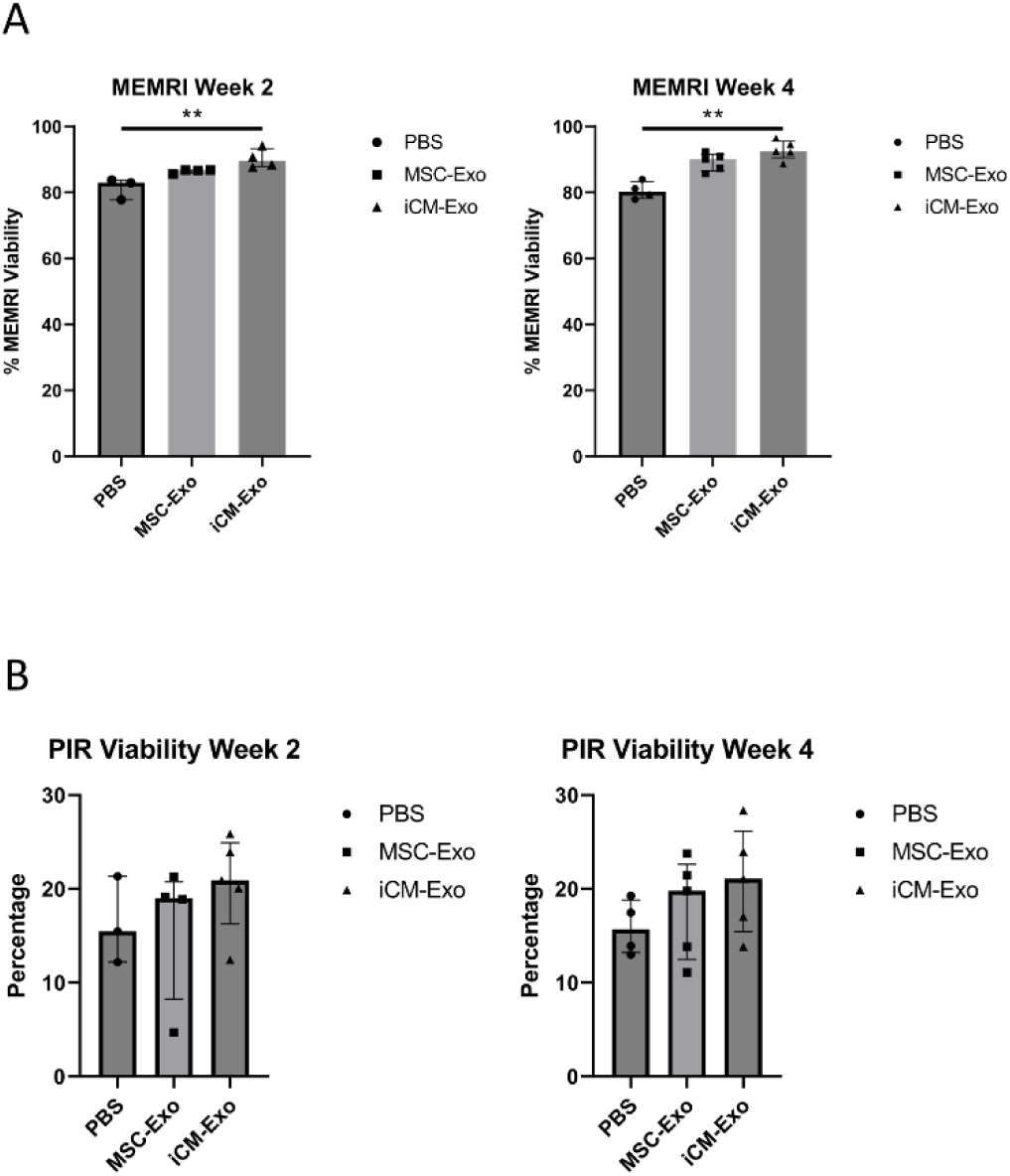
Cardiac MRI of porcine hearts treated with PBS, MSC-Exo, or iCM-Exo. Quantification of (A) manganese enhanced magnetic resonance imaging percent viability and (B) peri-infarct region percent viability. Statistical significance was determined by Kruskal Wallis (iCM-Exo, n=5; MSC-Exo, n=5; PBS n=4) with multiple comparisons. Median with interquartile range is shown. Only significance between control and treatment groups within each week are shown. (**p<0.01) iCM-Exo: exosomes derived from induced pluripotent stem cell derived cardiomyocytes; MEMRI: manganese enhanced MRI; MRI: magnetic resonance imaging; MSC-Exo: exosomes derived from mesenchymal stem cells; PIR: peri infarct region

### Exosomes from iCMs and MSCs improved cardiac function through similar mechanisms of action using different molecular pathways

Differential gene analysis provided insight into the mechanism of action of the two exosome treatment groups to rescue the damaged tissue, contributing to improved peri-infarct region viability and LVEF. When comparing the iCM-Exo vs. PBS treated peri-infarct region, 9782 genes were co-expressed, 1560 were uniquely expressed in iCM-Exo vs. 693 in PBS group (Fig 3A). When comparing MSC-Exo vs. PBS treated peri-infarct region specimens, 10147 genes were co-expressed and 542 were uniquely expressed in the MSC-Exo vs. 328 in PBS group (Fig 3A). iCM-Exo and MSC-Exo treated peri-infarct region tissue shared 10033 genes with 1309 unique to iCM-Exo vs. 656 unique to MSC-Exo treatment groups (Fig 3A).

**Figure 3.**
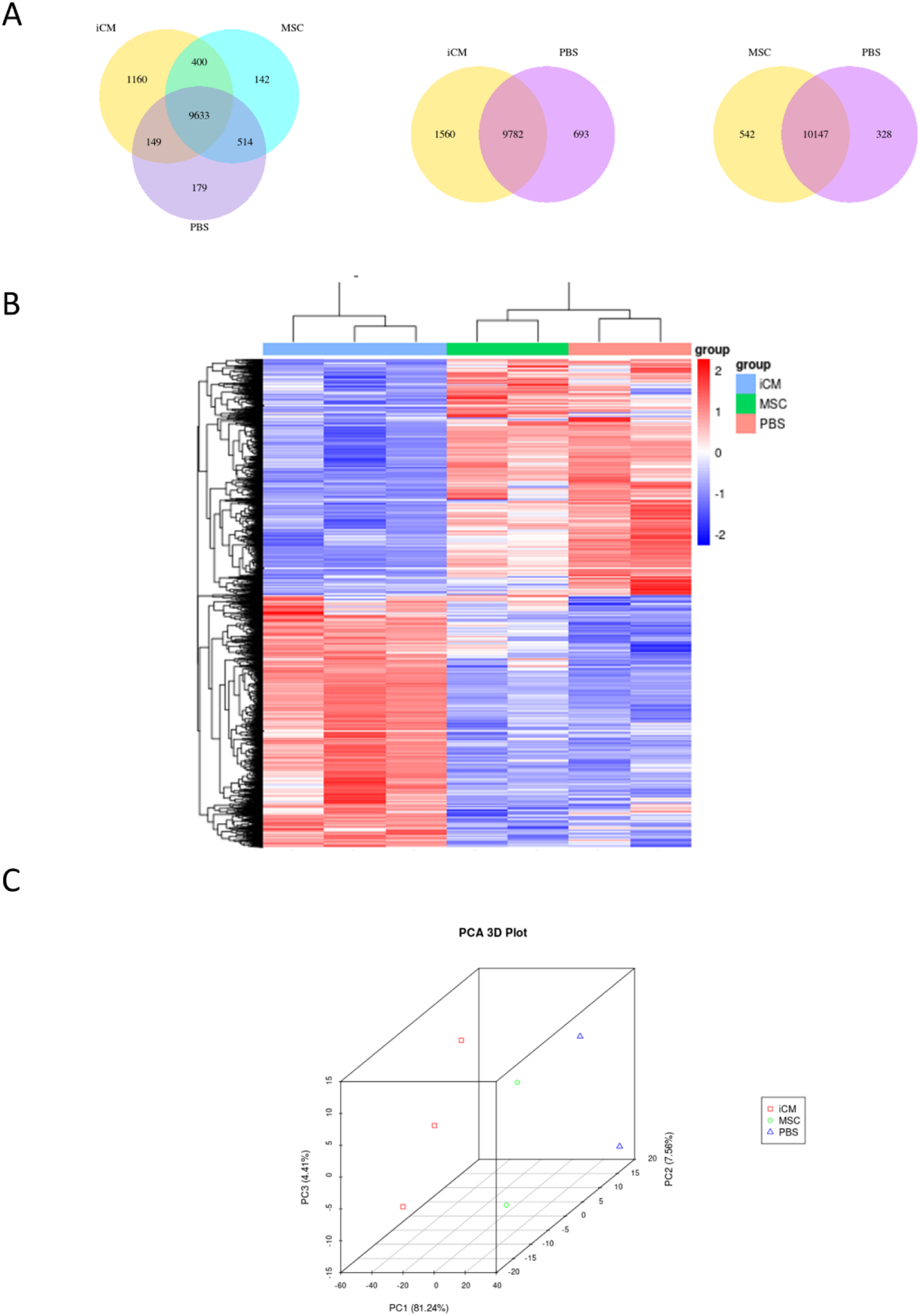
Next Generation Sequencing of RNA of treated peri-infarct region tissue. A) Venn diagram of unique and shared genes expressed in tissue treated with either PBS, iCM-Exosomes, or MSC-exosomes. B) Hierarchical clustering of differentially expressed genes between the three treatment groups. C) 3D PCA plot of each sample submitted for sequencing. iCM-Exo: exosomes derived from induced pluripotent stem cell derived cardiomyocytes; MSC-Exo: exosomes derived from mesenchymal stem cells; PIR: peri infarct region

Differential gene analysis demonstrated that iCM-Exo treated peri-infarct region tissue had 4453 significantly up-regulated gene expression and 4230 down-regulated expression when compared to PBS treated tissue (Fig 3B). On the other hand, MSC-Exo treatment only had 185 up-regulated genes and 272 down-regulated genes when compared to PBS treated tissue (Fig 3B). Both treatment groups demonstrated high differential expression of genes associated with PI3K/Akt/mTOR signaling pathway and focal adhesion, suggesting that both iCM-Exo and MSC-Exo are promoting rescue of damaged cells, possibly by proliferation^24–26^. Each treatment group was also clustered together, and variation observed in the PCA plot may be due to individual difference (Fig 3C).

Kyoto Encyclopedia of Genes and Genomes (KEGG) pathway analysis and Gene Ontology (GO) pathway analysis of differentially expressed genes (DEGs) showed that iCM-Exo and MSC-Exo both improved cardiac function through similar mechanisms of action. Notably, GO analysis of iCM-Exo vs. MSC-Exo treated hearts displayed higher expression of inflammation related genes in iCM-Exo, including but not limited to cytokines and macrophages (Fig 4A). Both treatments had high differential expression of genes associated with upregulation in PI3K/Akt/mTOR signaling pathway, focal adhesion, and other notable pathways associated with cellular proliferation, suggesting that both iCM-Exo and MSC-Exo are promoting rescue of damaged cells compared to PBS treatment (Fig 4B and 4C). In addition to the PI3K Akt mTOR and focal adhesion pathways, top targets of up-regulated DEGs in iCM-Exo vs PBS showed alterations in Yes-associated protein (YAP) and Hippo (Fig 4B). Similarly, upregulated DEGs in MSC-Exo vs PBS show changes in YAP1 and TGF-β (Fig 4C).

**Figure 4.**
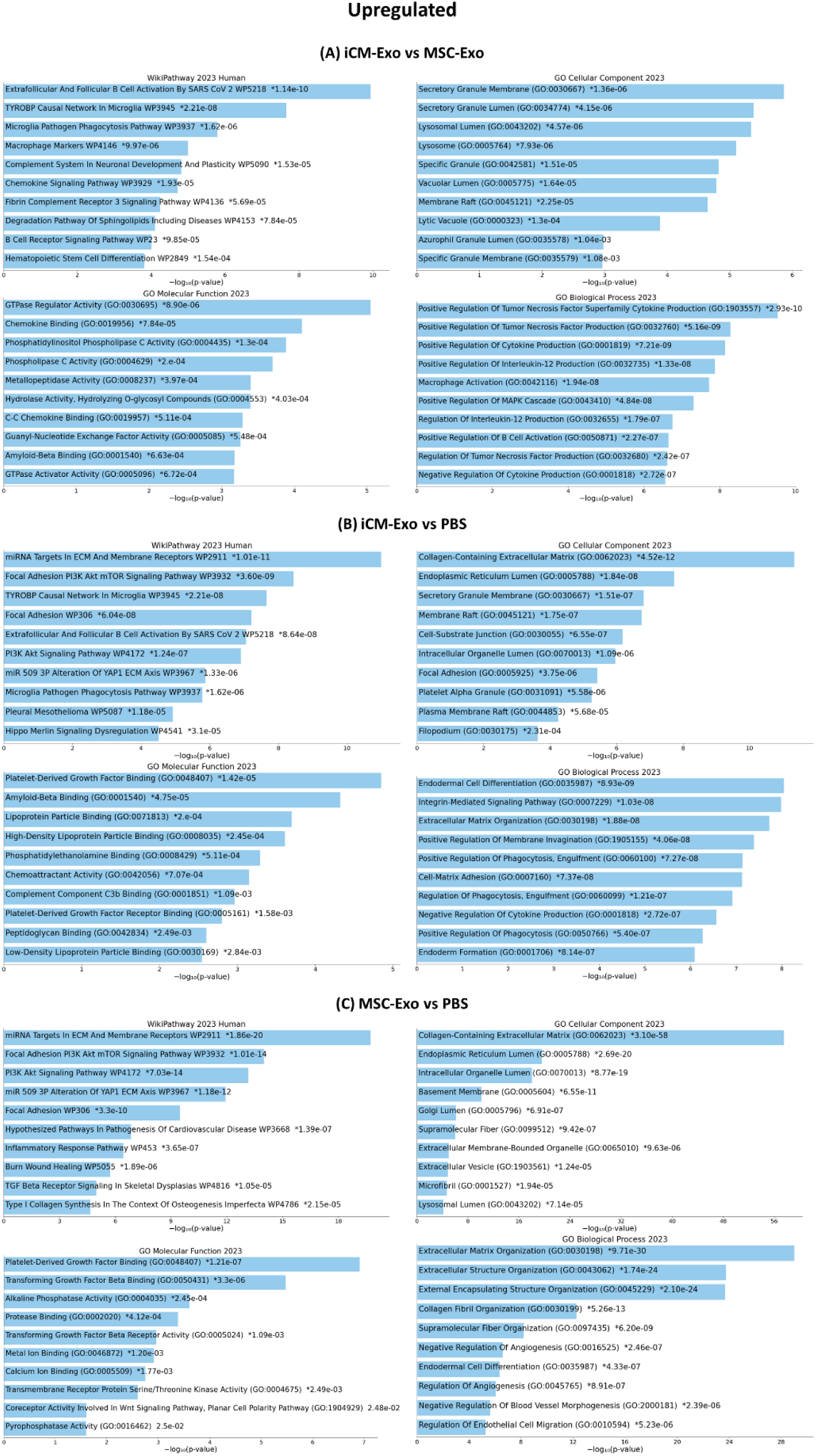
Upregulated differentially expressed genes from treated peri-infarct tissue. Kyoto Encyclopedia of Genes and Genomes Pathway Analysis and Gene Ontology analysis of A) Upregulated differentially expressed analysis in iCM-Exo vs MSC-Exo treated tissue. B) Upregulated differentially expressed analysis in iCM-Exo vs PBS treated tissue. C) Upregulated differentially expressed analysis in MSC-Exo vs PBS treated tissue. Graphs were created and obtained from Enrichr. iCM-Exo: exosomes derived from induced pluripotent stem cell derived cardiomyocytes; MSC-Exo: exosomes derived from mesenchymal stem cells

Both treatment groups also downregulated mitochondrial components related to oxidative phosphorylation and electron transport chain, which may facilitate hibernation and preservation of the damaged myocardium (Fig 5A, B, C). iCM-Exo vs. MSC-Exo treatment showed significant down-regulation in mitochondrial components, suggesting an enhanced suppression of mitochondrial energetics in iCM-Exo treatment groups compared to MSC-Exo (Fig5A). Both GO and KEGG pathway analyses, comparing iCM-Exo and MSC-Exo to PBS, showed down-regulation in mitochondrial function (Fig 5B, C).

**Figure 5.**
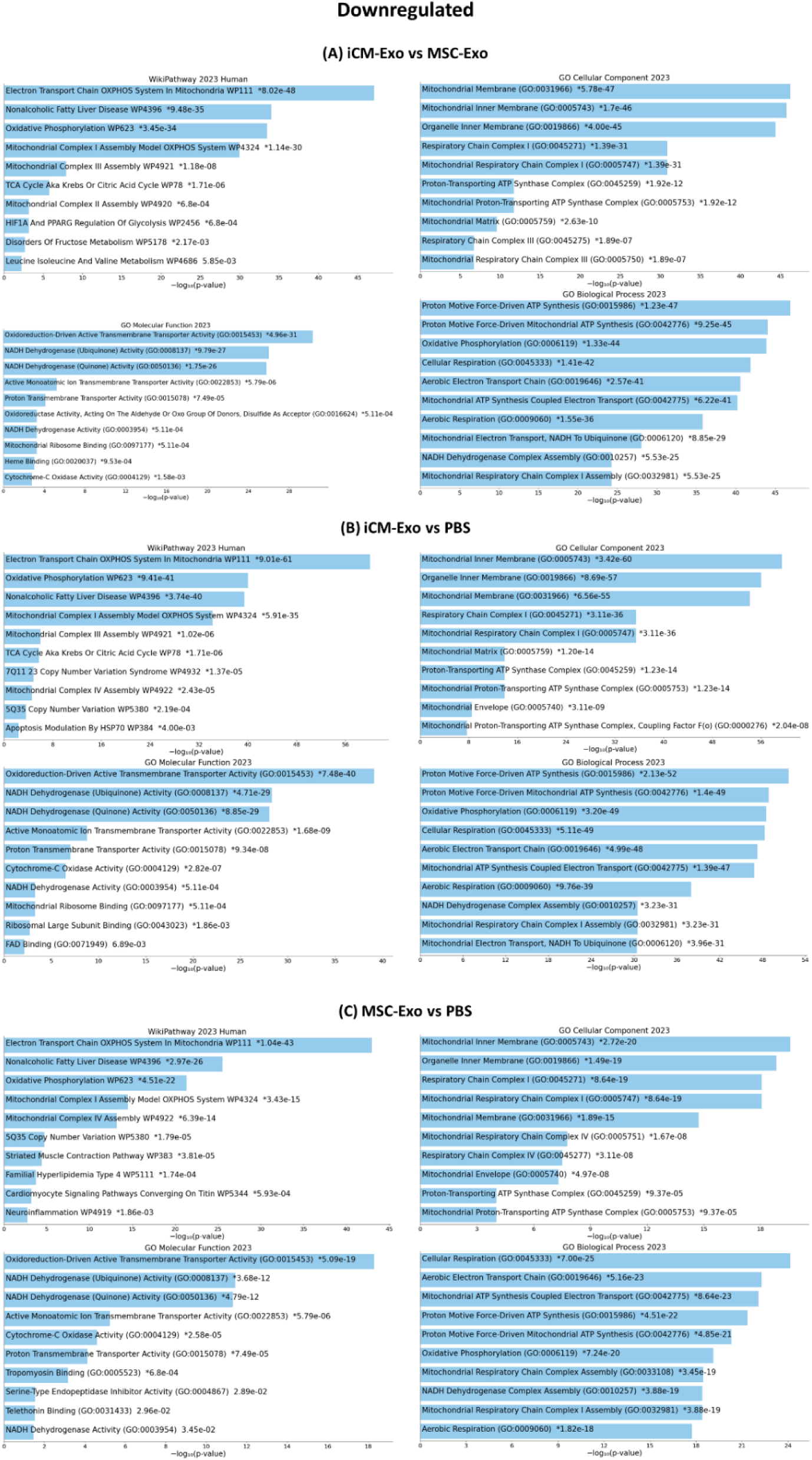
Downregulated differentially expressed genes from treated peri-infarct tissue. Kyoto Encyclopedia of Genes and Genomes Pathway Analysis and Gene Ontology analysis of A) Downregulated differentially expressed analysis in iCM-Exo vs MSC-Exo treated tissue. B) Downregulated differentially expressed analysis in iCM-Exo vs PBS treated tissue. C) Downregulated differentially expressed analysis in MSC-Exo vs PBS treated tissue. Graphs were created and obtained from Enrichr. iCM-Exo: exosomes derived from induced pluripotent stem cell derived cardiomyocytes; MSC-Exo: exosomes derived from mesenchymal stem cells; PIR: peri infarct region

### RNA Next Generation Sequencing of MSC-Exo or iCM-Exo demonstrate that their mechanism of actions is derived from different molecular payload

Following transcriptomic analysis of the porcine peri-infarct region tissue, the contents of MSC-Exo vs. iCM-Exo were investigated by differential gene expression analysis by exosomal Next Generation Sequencing. There were distinct groups of significantly expressed miRNA population between MSC-Exo vs. iCM-Exo (Fig 6). Although the porcine transcriptomic data suggest similar mechanisms of action in restoring cardiac function, MSC-Exo vs. iCM-Exo do so through different pathways and molecules. Of note, there appears to be numerous differentially expressed miRNAs in both exosomes related to the up-regulation of proliferation and down-regulation of mitochondrial components seen in treated peri-infarct region. Analysis of mRNAs in MSC-Exo and iCM-Exo suggests a similar trend in which proliferation and mitochondrial components are affected (Fig 7). Further studies are required to make any conclusions on how these miRNAs and mRNAs affect proliferation and mitochondrial function *in vivo* and to identify the molecular effectors of cardiac restoration.

**Figure 6.**
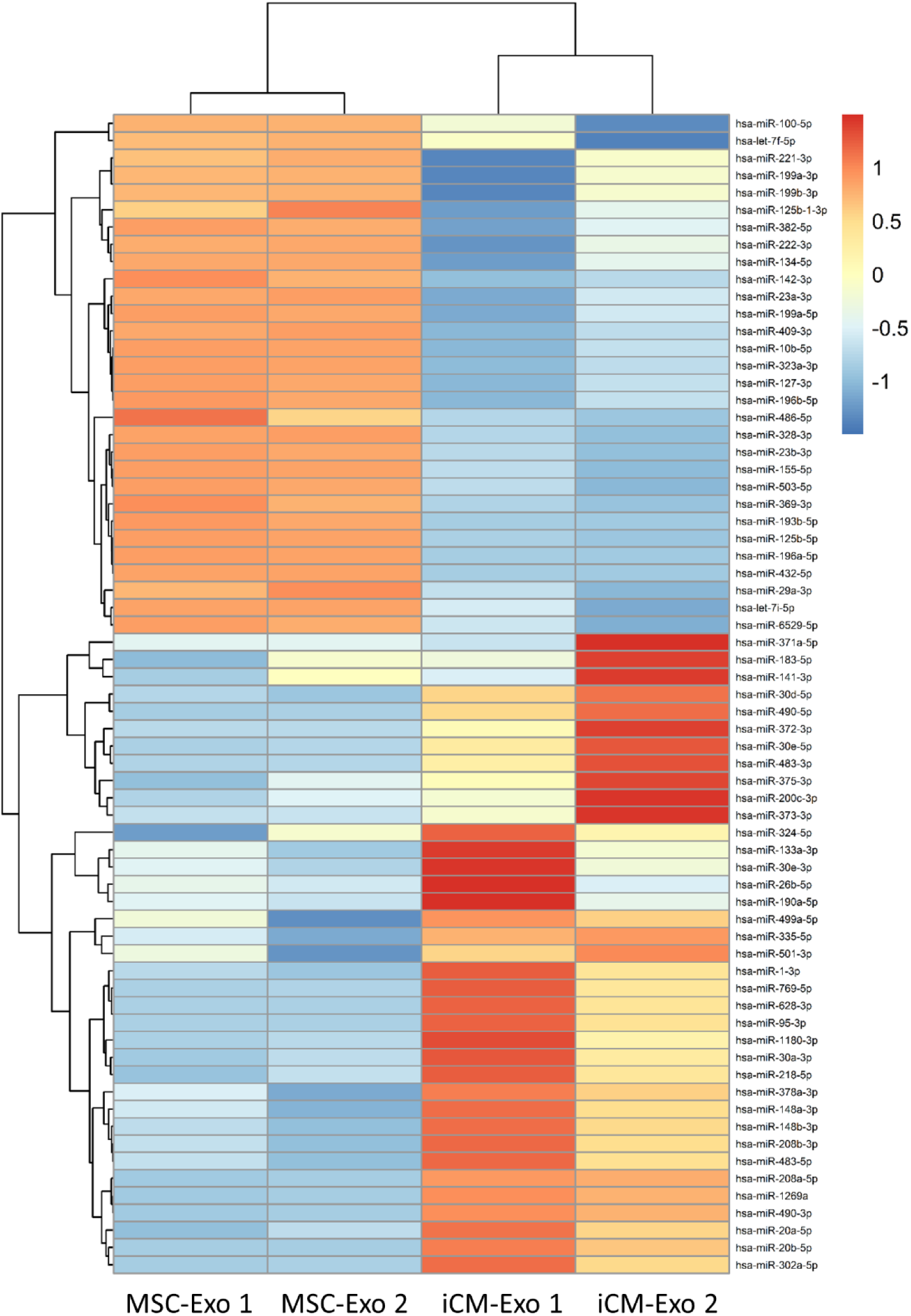
Heat map of significantly differentially expressed miRNA in MSC-Exo and iCM-Exo. Significantly differentially expressed miRNA were determined from MSC-Exo and iCM-Exo (n=2 each) by Next Generation Sequencing, demonstrating different populations of miRNAs produced in exosomes from different origin cells.

**Figure 7.**
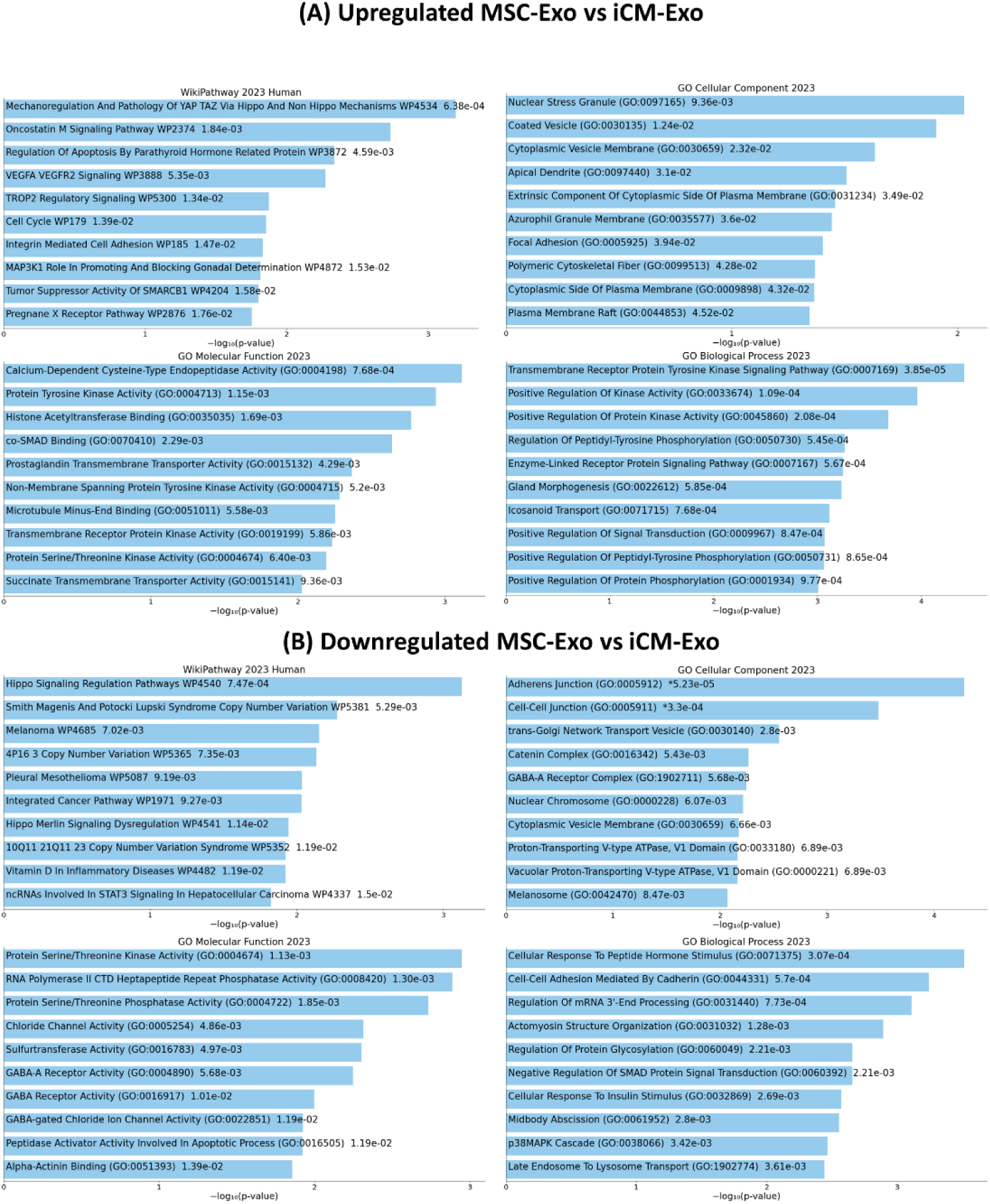
Differentially expressed genes from MSC-Exo and iCM-Exo content. Kyoto Encyclopedia of Genes and Genomes Pathway Analysis and Gene Ontology analysis of A) Upregulated differentially expressed analysis in MSC-Exo vs iCM-Exo. B) Downregulated differentially expressed analysis in MSC-Exo vs iCM-Exo. Graphs were created and obtained from Enrichr. iCM-Exo: exosomes derived from induced pluripotent stem cell derived cardiomyocytes; MSC-Exo: exosomes derived from mesenchymal stem cells

## Discussion

In this study, we demonstrated significant improvement in cardiac function and morphology of the injured porcine myocardium with iCM-Exo and a trend towards improvement with MSC-Exo treatments. Manganese enhanced magnetic resonance imaging demonstrated that iCM-Exo treated hearts had significant increase in viable myocardium at both weeks 2 and 4 while delayed gadolinium enhancement magnetic resonance imaging showed that MSC-Exo significantly decreased the scar size at week 4 when compared to the control heart. The marked improvements seen in cardiac function are conferred by miRNA carried by the exosomes, particularly by cardioprotective reduction in metabolism during acute myocardial injury while promoting concurrent cardiomyocyte cell cycle re-entry and proliferation. Exosome treatments improved the heart function and morphology by increasing the viable myocardium in peri-infarct region, resulting in sustained recovery over the 4-week period. The PBS treated control hearts deteriorated significantly over time when compared to the exosome treated hearts. Tissue from the peri-infarct region in both exosome treated hearts displayed an increase in PI3K Akt signaling pathway and focal adhesion and a reduction in mitochondrial components. There was no noted up-regulation of autophagy/mitophagy markers. However, this could be due to the harvesting of tissue 4 weeks after administering exosome treatments. Though the exosomes from the two parent cells contained different payloads, the overall effect they elicited on the recipient tissue was notably similar.

There are numerous differentially expressed miRNAs in both exosome groups related to proliferation and mitochondrial energetics. Several notable miRNAs, which target cellular proliferation, include hsa-miR-1-3p and the let-7 family. hsa-miR-1-3p is known to target the Notch pathway and is known to be a key player in regulating cardiac and skeletal muscle development^27–29^. Furthermore, it has also been shown that miR-1 improves heart function and drives cardiac differentiation in an infarcted murine model through the PTEN/Akt pathway^30^, which is consistent with our results discussed above in pig transcriptomic analysis. hsa-miR-199a-5p has also been found to promote cell proliferation^31–32^, and hsa-miR-302a-5p is part of the miR302-367 cluster, where one of the targets for cardiac repair is the Hippo signaling pathway^33^. mRNA transcriptomic analysis of iCM-Exo and MSC-Exo also suggested a similar trend where proliferation is a candidate for the mechanism of action. Hippo and YAP genes were upregulated in both iCM-Exo and MSC-Exo mRNA populations. Of note, hsa-miR-20b was found to be upregulated in iCM-Exo and downregulated in MSC-Exo, which we previously found to be part of the miR-106a-363 cluster of iCM-Exo that restores injured myocardium through the Notch3 pathway^11^.

There are also differentially expressed miRNAs in both exosome groups related to mitochondria bioenergetics. Of note, the miRNAs, which activate cellular proliferation, also target mitochondrial bioenergetics to create a complementary network of biological processes to rest and restore the acutely injured myocardium. One example of overlap is hsa-miR-1-3p and the let-7 family. hsa-miR-1-3p is activated during myogenesis to enter the mitochondria and enhance translation of bioenergetic proteins^34–35^. The let-7 family and hsa-miR-302a-5p also have many putative targets mapping to the mitochondrial genome^34–36^. Furthermore, hsa-miR-199a-5p has been found to target mitochondrial biogenesis through PGC-1α^31–32^. Electron transport chain complexes of reperfusion-injured mitochondria produce reactive oxygen species and initiate apoptotic processes, compromising the function of neighboring tissue. We speculate that the downregulation of oxidative phosphorylation components in the peri-infarct region prevents the generation of pro-apoptotic effectors in the acutely damaged myocardium. Reduced metabolic function also suggests hibernation of the injured myocardium for tissue homeostasis and self-protection, which enables longitudinal recovery of myocardial function through cellular proliferation.

Decreased metabolism and hibernating myocardium were proposed several decades ago and have been validated clinically^37–39^. This study is one of the first demonstrations of treatment contributing to the homeostatic effects in the infarcted heart where the downregulation of mitochondrial components contributed to the hibernating myocardium and reduced metabolism as a protective measure for the injured heart. Specifically, our study confirms the effect of up-regulation of mTOR signaling in reducing the metabolism of treated porcine peri-infarct region tissue. Other studies have confirmed this finding by showing that the downregulation or deletion of mTOR signaling can regulate cardiac metabolism by increasing glucose oxidation while decreasing fatty acid oxidation^40^. We have also previously demonstrated that iCM-Exo treatments in murine injury model promoted autophagy for myocardial repair through down-regulation of mTOR^9^, suggesting that the different sources of exosomes may contribute to modulate multiple facets of cardiac bioenergetics to restore and improve cardiac function. Our data confirmed the reported role of mTOR in altering cardiomyocyte metabolism and cell proliferation^41–42^.

The study data offers evidence that proliferation is a strong candidate for mechanism of action of iCM-Exo and MSC-Exo in improving cardiac function. Top hits in up-regulated DEGs in iCM-Exo vs. PBS showed significant difference in YAP and Hippo (Fig 4B) and up-regulated DEGs in MSC-Exo vs PBS showed changes in YAP1 and TGF-β (Fig 4C). Hippo signaling and PI3K-Akt signaling are both pathways implicated in cardiac proliferation while focal adhesion is a less established pathway related to cardiac regeneration^24, 25, 43–44^. YAP is a positive regulator for cardiac cell proliferation, acting as a downstream, terminal effector in the Hippo signaling pathway^25, 43, 45^. YAP has also been shown to stimulate cell cycle re-entry and adult cardiomyocyte proliferation in a murine myocardial infarction model^46^. Hippo and TGF-β are known to regulate the adult cardiomyocyte cell-cycle. Specifically, TGF-β superfamily plays major roles in heart development and interacts with other signaling pathways, contributing to cardiac proliferation^47^. The dual mechanism of cell proliferation and myocardial hibernation represents the necessary homeostatic process to restore and improve cardiac function. Transcriptomic analysis of MSC-Exo and iCM-Exo content shows distinct populations of miRNA, suggesting that exosomes from two different parent cells were contributing to a similar mechanism of action with different pathways and molecules.

Based on the dual mechanism of action, our data suggest that the partially injured but viable cardiomyocytes are rescued through cell-cycle re-entry or proliferation and severely damaged cells undergo reduced metabolics and hibernation to recover function. However, this requires further study not only on how different cell types react to exosome treatments but also to what degree exosome contents can affect the cells. Furthermore, while numerous components of exosome content, such as miRNA, have been studied extensively, there are also many miRNA and other cellular components with little known function in the context of complex cardiovascular homeostasis.

## Conclusion

In an acute porcine myocardial infarction model, we investigated the effect of MSC-Exo and iCM-Exo treatments on injured myocardium. Both iCM-Exo and MSC-Exo treatment can improve the cardiac function and morphology of injured myocardium. Comparatively, PBS treated control hearts deteriorated significantly overtime. Transcriptomic analysis from the peri-infarct region in both MSC-Exo and iCM-Exo treated hearts showed a significant reduction in myocardial metabolism and an increase in proliferation signal pathways. Next Generation Sequencing of exosome cargo of MSC-Exo and iCM-Exo showed different payloads but elicited similar overall effect on treated tissue.

### What is known?

- Heart failure is the leading cause of hospital admission in the United States with continued increase in patient population.
- Transplantation of mesenchymal stem cells or induced pluripotent stem cell derived cardiomyocytes improve heart function after myocardial infarction in pre-clinical models.
- Exosomes regulate the underlying paracrine mechanism of action of stem cells and contain a wide-range of payload, containing diverse set of molecular, cellular and proteomic components.

### What new information does this article contribute?

- Exosomes from both MSCs and iCMs were shown to improve cardiac function in an acute porcine myocardial injury model.
- A dual mechanism of action was delineated from our multi-omic investigation of *ex vivo* porcine myocardium in both treatment groups.
- The dual mechanism consists of the following: 1) cardioprotective reduction in metabolism and 2) promoted cardiomyocyte cell cycle re-entry and proliferation

### Novelty and significance

In an acute porcine myocardial infarction model, we demonstrated that the exosomes from both MSCs and iCMs improve the cardiac function of injured myocardium. We further delineated a dual mechanism of action from our multi-omic investigation of *ex vivo* porcine myocardium in both treatment groups. Through transcriptomic analysis from the peri-infarct region in treated heart, we demonstrated that both MSC-Exo and iCM-Exo confer improved cardiac function through a significant reduction in myocardial metabolism and an increase in proliferation signal pathways. Together, this investigation provides a pre-clinical validation of a novel biologic to treat heart failure. Further investigation of optimal dose-response and administration frequency as well as the effect in a chronic porcine heart failure model need to be elucidated for comprehensive translational research.

## Non-standard Abbreviations and Acronyms

MSC: human bone marrow derived mesenchymal stem cells
iCM: induced pluripotent stem cell derived cardiomyocytes
MSC-Exo: exosomes derived from mesenchymal stem cells
iCM-Exo: exosomes derived from induced pluripotent stem cell-derived cardiomyocytes
LVEF: left ventricular ejection fraction
LVESV: left ventricular end-systolic volume
LVEDV: left ventricular end-diastolic volume
KEGG: Kyoto Encyclopedia of Genes and Genomes
GO: Gene Ontology
DEG: differentially expressed genes

## Acknowledgements

None.

## References

1. Amado LC, Saliaris AP, Schuleri KH, John MS, Xie J-S, Cattaneo S, Durand DJ, Fitton T, Kuang JQ, Stewart G, et al. Cardiac repair with intramyocardial injection of allogeneic mesenchymal stem cells after myocardial infarction. Proc Natl Acad Sci U S A. 2005;102:11474–11479.

2. Armiñán A, Gandía C, García-Verdugo JM, Lledó E, Trigueros C, Ruiz-Saurí A, Miñana MD, Solves P, Payá R, Montero JA, et al. Mesenchymal stem cells provide better results than hematopoietic precursors for the treatment of myocardial infarction. J Am Coll Cardiol. 2010;55:2244–2253.

3. Rojas SV, Kensah G, Rotaermel A, Baraki H, Kutschka I, Zweigerdt R, Martin U, Haverich A, Gruh I, Martens A. Transplantation of purified iPSC-derived cardiomyocytes in myocardial infarction. PloS One. 2017;12:e0173222.

4. Mozaffarian D, Benjamin EJ, Go AS, Arnett DK, Blaha MJ, Cushman M, de Ferranti S, Després JP, Fullerton HJ, Howard VJ, et al. Heart disease and stroke statistics—2015 update. Circulation. 2015;131:e29–e322.

5. Shiba Y, Gomibuchi T, Seto T, Wada Y, Ichimura H, Tanaka Y, Ogasawara T, Okada K, Shiba N, Sakamoto K, et al. Allogeneic transplantation of iPS cell-derived cardiomyocytes regenerates primate hearts. Nature. 2016;538:388–391.

6. Zhao M, Nakada Y, Wei Y, Bian W, Chu Y, Borovjagin AV, Xie M, Zhu W, Nguyen T, Zhou Y, et al. Cyclin D2 overexpression enhances the efficacy of human induced pluripotent stem cell–derived cardiomyocytes for myocardial repair in a swine model of myocardial infarction. Circulation. 2021;144: 210–228.

7. Gao L, Wang L, Wei Y, Krishnamurthy P, Walcott GP, Menasché P, Zhang J. Exosomes secreted by hiPSC-derived cardiac cells improve recovery from myocardial infarction in swine. Sci Transl Med. 2020;12:eaay1318.

8. Yang PC. Induced pluripotent stem cell (iPSC)–derived exosomes for precision medicine in heart failure. Circ Res. 2018;122:661–663.

9. Santoso MR, Ikeda G, Tada Y, Jung JH, Vaskova E, Sierra RG, Gati C, Goldstone AB, von Bornstaedt D, Shukla P, et al. Exosomes from induced pluripotent stem cell–derived cardiomyocytes promote autophagy for myocardial repair. J Am Heart Assoc. 2020;9:e014345.

10. Ikeda G, Santoso MR, Tada Y, Li AM, Vaskova E, Jung JH, O’Brien C, Egan E, Ye J, Yang PC. Mitochondria-rich extracellular vesicles from autologous stem cell–derived cardiomyocytes restore energetics of ischemic myocardium. J Am Coll Cardiol. 2021;77:1073–1088.

11. Jung JH, Ikeda G, Tada Y, von Bornstädt D, Santoso MR, Wahlquist C, Rhee S, Jeon YJ, Yu AC, O’brien CG, et al. miR-106a–363 cluster in extracellular vesicles promotes endogenous myocardial repair via Notch3 pathway in ischemic heart injury. Basic Res Cardiol. 2021;116:19.

12. Davidson SM, Riquelme JA, Zheng Y, Vicencio JM, Lavandero S, Yellon DM. Endothelial cells release cardioprotective exosomes that may contribute to ischaemic preconditioning. Sci Rep. 2018;8:15885.

13. Vicencio JM, Yellon DM, Sivaraman V, Das D, Boi-Doku C, Arjun S, Zheng Y, Riquelme JA, Kearney J, Sharma V, et al. Plasma exosomes protect the myocardium from ischemia-reperfusion injury. J Am Coll Cardiol. 2015;65:1525–1536.

14. Minghua W, Zhijian G, Chahua H, Qiang L, Minxuan X, Luqiao W, Weifang Z, Peng L, Biming Z, Lingling Y, et al. Plasma exosomes induced by remote ischaemic preconditioning attenuate myocardial ischaemia/reperfusion injury by transferring miR-24. Cell Death Dis. 2018;9:320.

15. Li J, Rohailla S, Gelber N, Rutka J, Sabah N, Gladstone RA, Wei C, Hu P, Kharbanda RK, Redington AN. microRNA-144 is a circulating effector of remote ischemic preconditioning. Basic Res Cardiol. 2014;109:423.

16. Zhao JF, Rodger CE, Allen GFG, Weidlich S, Ganley IG. HIF1α-dependent mitophagy facilitates cardiomyoblast differentiation. Cell Stress. 2020;4:99–113.

17. Honka H, Solis-Herrera C, Triplitt C, Norton L, Butler J, DeFronzo RA. Therapeutic manipulation of myocardial metabolism: JACC state-of-the-art review. J Am Coll Cardiol. 2021;77:2022–2039.

18. Tada Y, Heidary S, Tachibana A, Tachibana A, Zaman J, Neofytou E, Dash R, Wu JC, Yang PC. Myocardial viability of the peri-infarct region measured by T1 mapping post manganese-enhanced MRI correlates with LV dysfunction. Int J Cardiol. 2019;281:8–14.

19. Dash R, Kim PJ, Matsuura Y, Ikeno F, Metzler S, Huang NF, Lyons JK, Nguyen PK, Ge X, Foo CWP, et al. Manganese-enhanced magnetic resonance imaging enables in vivo confirmation of peri-infarct restoration following stem cell therapy in a porcine ischemia– reperfusion model. J Am Heart Assoc. 2015;4:e002044.

20. Chen EY, Tan CM, Kou Y, Duan Q, Wang Z, Meirelles GV, Clark NR, Ma’ayan A. Enrichr: interactive and collaborative HTML5 gene list enrichment analysis tool. BMC Bioinformatics. 2013;14:128.

21. Kuleshov MV, Jones MR, Rouillard AD, Fernandez NF, Duan Q, Wang Z, Koplev S, Jenkins SL, Jagodnik KM, Lachmann A, et al. Enrichr: a comprehensive gene set enrichment analysis web server 2016 update. Nucleic Acids Res. 2016;44:W90–W97.

22. Xie Z, Bailey A, Kuleshov MV, Clarke DJB., Evangelista JE, Jenkins SL, Lachmann A, Wojciechowicz ML, Kropiwnicki E, Jagodnik KM, et al. Gene set knowledge discovery with Enrichr. Curr Protoc. 2021;1:e90.

23. Clarke DJB, Jeon M, Stein DJ, Moiseyev N, Kropiwnicki E, Dai C, Xie Z, Wojciechowicz ML, Litz S, Hom J, et al. Appyters: turning Jupyter Notebooks into data-driven web apps. Patterns. 2021;2:100213.

24. Afify AY. A miRNA’s insight into the regenerating heart: a concise descriptive analysis. Heart Fail Rev. 2020;25:1047–1061.

25. Hashmi S, Ahmad HR. Molecular switch model for cardiomyocyte proliferation. Cell Regen. 2019;8:12–20.

26. Lin Z, Zhou P, von Gise A, Gu F, Ma Q, Chen J, Guo H, van Gorp PR, Wang DZ, Pu WT. Pi3kcb links Hippo-YAP and PI3K-AKT signaling pathways to promote cardiomyocyte proliferation and survival. Circ Res. 2015;116:35–45.

27. Kwon C, Han Z, Olson EN, Srivastava D. microRNA1 influences cardiac differentiation in Drosophila and regulates Notch signaling. Proc Natl Acad Sci U S A. 2005;102:18986– 18991.

28. Ivey KN, Muth A, Arnold J, King FW, Yeh RF, Fish JE, Hsiao EC, Schwartz RJ, Conklin BR, Bernstein HS, et al. microRNA regulation of cell lineages in mouse and human embryonic stem cells. Cell Stem Cell. 2008;2:219–229.

29. Zhao Y, Samal E, Srivastava D. Serum response factor regulates a muscle-specific microRNA that targets Hand2 during cardiogenesis. Nature. 2005;436:214–220.

30. Glass C, Singla DK. microRNA-1 transfected embryonic stem cells enhance cardiac myocyte differentiation and inhibit apoptosis by modulating the PTEN/Akt pathway in the infarcted heart. Am J Physiol Heart Circ Physiol. 2011;301:H2038–H2049.

31. Li Z, Liu L, Hou N, Song Y, An X, Zhang Y, Yang X, Wang J. miR-199-sponge transgenic mice develop physiological cardiac hypertrophy. Cardiovasc Res. 2016;110:258–267.

32. Chen HY, Lu J, Wang ZK, Yang J, Ling X, Zhu P, Zheng SY. hsa-miR-199a-5p protect cell injury in hypoxia induces myocardial cells via targeting HIF1α. Mol Biotechnol. 2022;64:482–492.

33. Tian Y, Liu Y, Wang T, Zhou N, Kong J, Chen L, Snitow M, Morley M, Li D, Petrenko N, et al. A microRNA-Hippo pathway that promotes cardiomyocyte proliferation and cardiac regeneration in mice. Sci Transl Med. 2015;7:279ra38-279ra38.

34. Borralho PM, Rodrigues CMP, Steer CJ. microRNAs in mitochondria: an unexplored niche. In: Santulli G, editor. microRNA: Basic Science. Advances in Experimental Medicine and Biology. New York City, NY: Springer International Publishing, Cham, 2015: 31–51.

35. Zhang X, Zuo X, Yang B, Li Z, Xue Y, Zhou Y, Huang J, Zhao X, Zhou J, Yan Y, et al. microRNA directly enhances mitochondrial translation during muscle differentiation. Cell. 2014;158:607–619.

36. Barrey E, Saint-Auret G, Bonnamy B, Damas D, Boyer O, Gidrol X. Pre-microRNA and mature microRNA in human mitochondria. PloS One. 2011;6:e20220.

37. Wijns W, Vatner SF, Camici PG. Hibernating myocardium. N Engl J Med. 1998;339:173– 181.

38. Schinkel AFL, Bax JJ, Poldermans D, Elhendy A, Ferrari R, Rahimtoola SH. Hibernating myocardium: diagnosis and patient outcomes. Curr Probl Cardiol. 2007;32:375–410.

39. Elsässer A, Schlepper M, Klövekorn WP, Cai WJ, Zimmermann R, Müller KD, Strasser R, Kostin S, Gagel C, Münkel B, et al. Hibernating myocardium: an incomplete adaptation to ischemia. Circulation. 1997;96:2920–2931.

40. Zhu Y, Soto J, Anderson B, Riehle C, Zhang YC, Wende AR, Jones D, McClain DA, Abel ED. Regulation of fatty acid metabolism by mTOR in adult murine hearts occurs independently of changes in PGC-1α. Am J Physiol Heart Circ Physiol. 2013;305:H41–H51.

41. Laplante M, Sabatini DM. Regulation of mTORC1 and its impact on gene expression at a glance. J Cell Sci. 2013;126:1713–1719.

42. Guertin DA, Stevens DM, Thoreen CC, Burds AA, Kalaany NY, Moffat J, Brown M, Fitzgerald KJ, Sabatini DM. Ablation in mice of the mTORC components raptor, rictor, or mLST8 reveals that mTORC2 is required for signaling to Akt-FOXO and PKCα, but not S6K1. Dev Cell. 2006;11:859–871.

43. von Gise A, Lin Z, Schlegelmilch K, Honor LB, Pan GM, Buck JN, Ma Q, Ishiwata T, Zhou B, Camargo FD, et al. YAP1, the nuclear target of Hippo signaling, stimulates heart growth through cardiomyocyte proliferation but not hypertrophy. Proc Natl Acad Sci U S A. 2012;109:2394–2399.

44. Xin M, Kim Y, Sutherland LB, Murakami M, Qi X, McAnally J, Porrello ER, Mahmoud AI, Tan W, Shelton JM, et al. Hippo pathway effector Yap promotes cardiac regeneration. Proc Natl Acad Sci U S A. 2013;110:13839–13844.

45. Boogerd CJ, Perini I, Kyriakopoulou E, Han SJ, La P, van der Swaan B, Berkhout JB, Versteeg D, Monshouwer-Kloots J, van Rooij E. Cardiomyocyte proliferation is suppressed by ARID1A-mediated YAP inhibition during cardiac maturation. Nat Commun. 2023;14:4716.

46. Lin Z, von Gise A, Zhou P, Gu F, Ma Q, Jiang J, Yau AL, Buck JN, Gouin KA, van Gorp PR, et al. Cardiac-specific YAP activation improves cardiac function and survival in an experimental murine MI model. Circ Res. 2014;115:354–363.

47. Sorensen DW, van Berlo JH. The role of TGF—β signaling in cardiomyocyte proliferation. Curr Heart Fail Rep. 2020;17:225–233.

